# Using LanM Enzymes to Modify Glucagon-Like Peptides 1 and 2 in *E.coli*

**DOI:** 10.1101/2023.03.20.532734

**Authors:** Camilla Kjeldgaard Larsen, Peter Lindquist, Mette Rosenkilde, Alice Ravn Madsen, Kim Haselmann, Tine Glendorf, Kjeld Olesen, Anne Louise Bank Kodal, Thomas Tørring

## Abstract

Selective modification of peptides is often exploited to improve pharmaceutically relevant properties of bioactive peptides like stability, circulation time, and potency. In Nature, natural products belonging to the class of ribosomally synthesized and post-translationally modified peptides (RiPPs) are known to install a number of highly attractive modifications with high selectivity. These modifications are installed by enzymes guided to the peptide by corresponding leader peptides removed as the last step of biosynthesis. Here, we exploit leader peptides and their matching enzymes to investigate the installment of D-Ala post-translationally in a critical position in the hormones, glucagon-like peptides (GLP) 1 and 2. We also offer insight into how precursor peptide design can modulate the modification pattern achieved.

## Introduction

Natural products remain a rich source of inspiration in the development of medications due to their notable structural and functional diversity and have driven our understanding of biology for the past century [1, 2]. <u>R</u>ibosomally synthesized and <u>P</u>ost-translationally modified <u>P</u>eptides (RiPPs) represent a large group of natural products. They are derived from ribosomally synthesized polypeptide precursors and despite their genetic simplicity, they are both structurally and functionally diverse [3, 4]. Their complex structure is installed through posttranslational modifications and endows the peptides with restricted conformational flexibility to allow better target recognition, increased metabolic and chemical stability, and improved chemical functionality [5]. From a medicinal perspective, these functionalities could improve peptide drugs with better shelf life, lower doses, and less frequent administration to patients [6].

One of the largest subclasses of RiPPs is the lanthipeptides characterized by thioether cross-links [7]. Lanthipeptides are grouped into five classes based on their biosynthetic machinery. In Class II lanthipeptides a multifunctional enzyme, LanM, catalyzes the posttranslational modifications of the genetically encoded precursor peptide (LanA) [8] by binding to the N-terminal part of the precursor peptide termed the leader sequence through a specific RiPP recognition element (RRE). Once bound to the LanM, the C-terminal core of the precursor peptide undergoes post-translational modifications to form dehydroalanines (Dha) and dehydrobutyrines (Dhb) and subsequently lanthionine (Lan), and methyllanthione (MeLan) moieties (Figure 1A). In this process, serine and threonine residues in the precursor peptide are processed in the dehydratase domain of the enzyme to Dha and Dhb, respectively, through a phosphorylation step using ATP and followed by an elimination with phosphate as the leaving group. Secondly, the cyclase domain of the enzyme is responsible for the installation of the thioether crosslinks formed by a Michael-type addition by the thiol of a Cys residue to the dehydrated amino acids [9–11]. In HalM2/HalA2, a representative of the class II lanthipeptides, HalM2 catalyzes seven phosphorylation, seven elimination, and four macrocyclizations - all chemoselective yielding one final product. While the details are still not completely understood, it is intriguing that the enzyme domains must cope with a substrate that changes drastically during maturation. Several recent papers have revealed that the enzyme is highly dynamic, and that binding of both leader – and core peptide lead to important structural changes that alter both reaction rates and fidelity [12, 13]. The leader sequence of the precursor peptide is enzymatically removed during biosynthetic maturation by dedicated proteases, and the modified core is released and transported out of the cell by transporters as the final RiPP [14]. Beyond the modifications installed by the leader-dependent enzymes, lanthipeptides are sometimes endowed with secondary modifications by tailoring enzymes. These enzymes can act on the mature peptide independently of the leader peptide, which will further increase the chemical diversity of the RiPP [4]. Some lanthipeptides are found with D-amino acids that are formed by hydrogenation of Dha and/or Dhb resulting in D-Ala and D-amino butyric acid, respectively. The enzymes that are responsible for the reduction reactions are called dehydrogenases and are collectively termed LanJ [15, 16]. The enzyme used in this study is called NpnJ, and is known to have a promiscuous substrate specificity in terms of peptide sequence, but specific to the reduction of Dha into D-Ala [17]. D-amino acids are of high interest, as they can improve properties like reduced susceptibility to proteolysis and induce turns [18–21]. As D-amino acids can not readly be incorporated recombinantly [22], D-amino acids containing peptides have been limited to chemical synthesis. A recent report from the Piel lab has demonstrated that a radical *S*-adenosyl methionine epimerase involved in proteusin biosynthesis can also be exploited to epimerize amino acids post-translationally [23].

**Figure 1.**
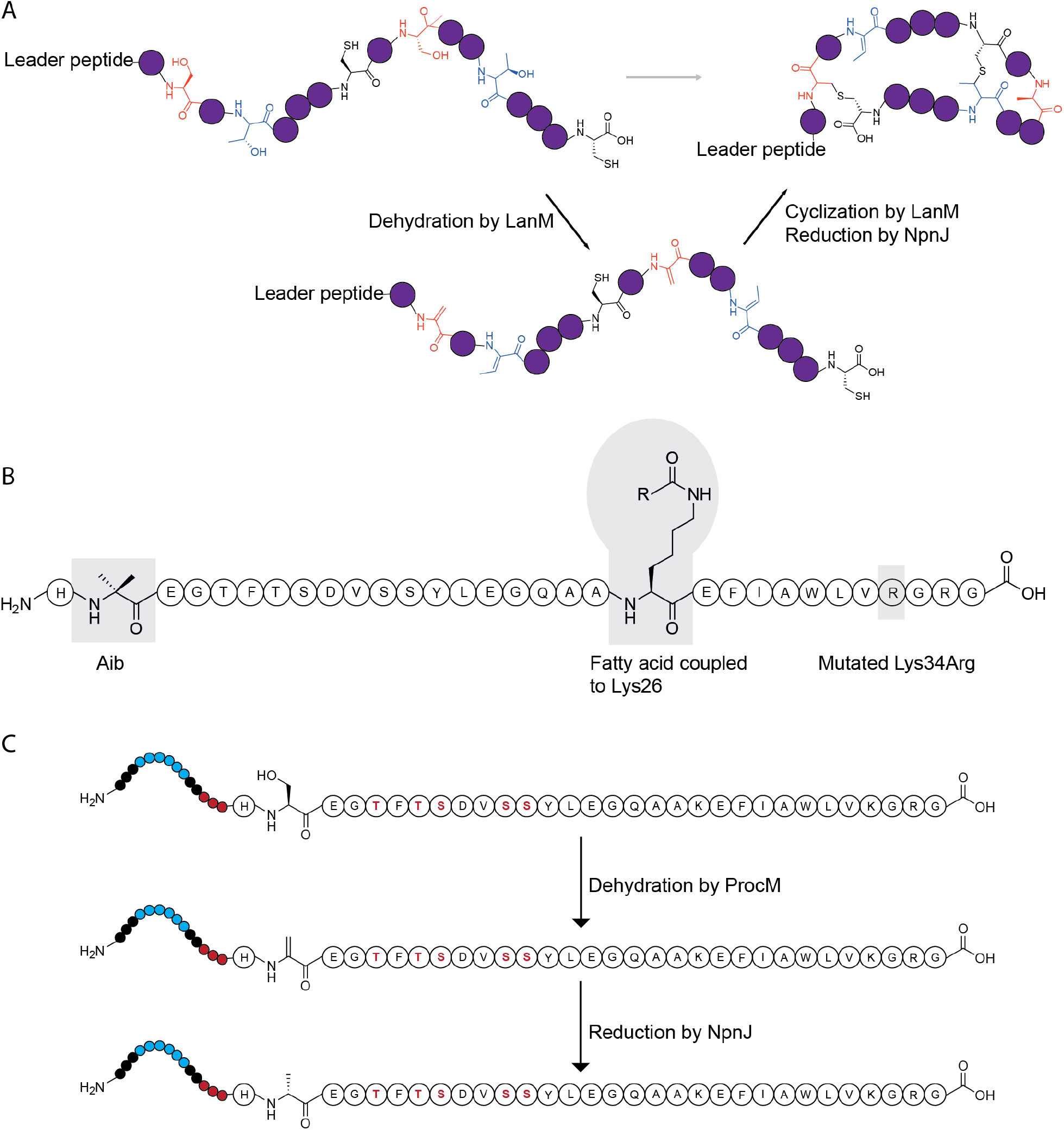
A) Hypothetical lanthipeptide and its post-translational modifications by LanM and NpnJ generation Dha, Dhb, Lan, MeLan, and D-Ala moieties. B) Sequences of semaglutide with highlighted changes, Aib, a fatty acid coupled to Lys26, and the mutated Lys34Arg. C) The recombinant biosynthesis of GLP-1 for the introduction of D-Ala8 by ProcM and NpnJ, additional Ser and Thr residues are highlighted.

Harnessing such modifications for use in biologically relevant peptides could assist the development of peptide based therapeutics. The 31-amino acid peptide hormone, glucagon-like peptide 1 (GLP-1), and the 33-amino acids glucagon-like peptide 2 (GLP-2) are both products of proglucagone derived from the glucagon gene [24]. GLP-1/-GLP-2 are secreted from intestinal endocrine L-cells located in the small and large intestines. GLP-1 is found in different variants, and the first predicted bioactive GLP-1 was composed of 37 amino acids (1-37). However, further studies revealed the six amino acids shorter bioactive version of GLP-1 (7-37) together with another one amino acid shorter GLP-1 (7-36)-amide, which is the result of enzymatic cleavage of the C-terminal glycine by peptidylglycine α-amidating monooxygenase (PAM) [25]. As an alternative to insulin-based therapies for people with type 2 diabetes, several modified peptide therapeutics based on GLP-1 have been approved one of them being semaglutide with an once weekly profile (Figure 1B). By chemically installing histidine and the non-proteogenic amino acid, amino-isobutyric acid (Aib) in positions 7-8, semaglutide is resistant towards proteolytic cleavage by dipeptidyl peptidase IV (DPP-IV). Adding a fatty acid chain to position 26 (Lys26) further increases the half-life in circulation by binding to serum albumin [26].

Here we show that it is possible to use class II lanthionine synthetases on GLP-1 and GLP-2 to selectively incorporate a D-Alanine on the second position to prevent proteolytic cleavage from DPP-IV (Figure 1C).

## Results and Discussion

### D-Ala could be an alternative to Aib in GLP-1 and GLP-2 analogues

The GLP-1 analogue, semaglutide contains the non-proteogenic amino acid 2-aminobutyric acid (Aib) in the eight position in the N-terminal to protect it from the endogenous DPP-IV. The two N-terminal amino acids His-Aib are introduced chemically which adds additional complexity to the production process. Inspired by Yang et al. [17], we hypothesized that enzymes known from RiPP biosynthesis could be exploited to introduce a D-Ala into this position and gain protection from DPP-IV while omitting a costly chemical step (Figure 2B, 2C) [27]. This relied on two assumptions: 1) that D-Ala would have a similar protection towards DPP-IV and 2) that we could use a combination of a dehydratase and the dehydrogenase, NpnJ, to install a D-Ala selectively on position 8 (GLP-1) and position 2 (GLP-2). GLP-2 was included as an additional peptide with sequence similarity to GLP-1.

**Figure 2.**
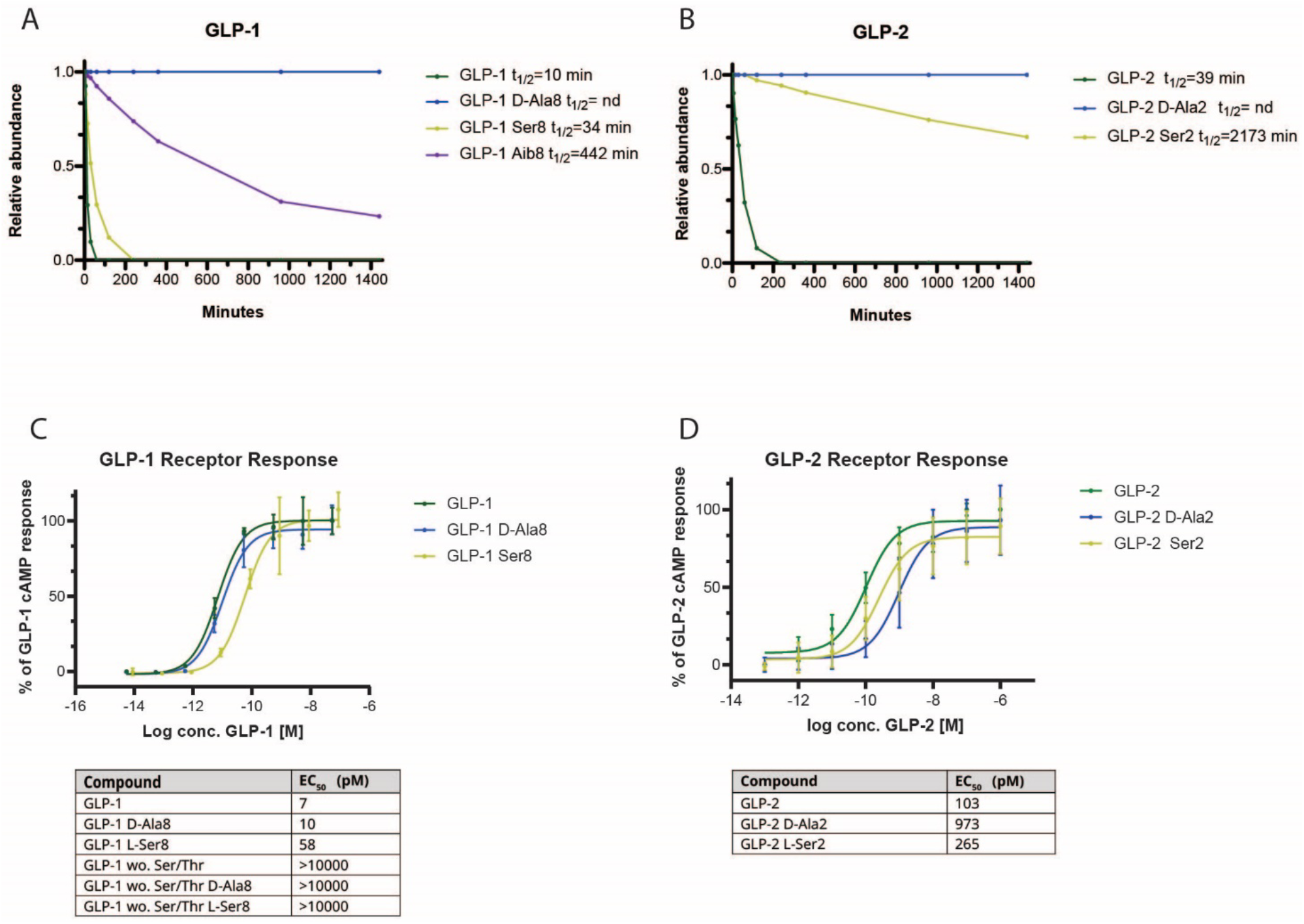
DPP-IV stability and potency of synthesized GLP-1 and GLP-2 variants.DPP-IV proteolysis assay of GLP-1 containing L-Ala, D-Ala, L-Ser, and Aib on residue 8. B) DPP-IV proteolysis assay of GLP-2 containing L-Ala, D-Ala and L-Ser on residue 2. C+D) Potency assays with GLP-1R and GLP-2R, respectively, made in duplicate.

### D-alanine prevents proteolytic degradation by DPP-IV

To address the first assumption, we synthesized a series of peptides using standard Fmoc-based peptide synthesis. We included the amino acids in semaglutide (Aib), natural GLP-1 (L-Ala), and our suggestion (D-Ala). All peptides were synthesized and purified in good yields and characterized using UHPLC-MS. Synthesized peptides with purity >86% were accepted. To investigate the stability in the presence of DPP-IV, synthesized peptides were mixed with DPP-IV and analyzed at different time points (Figure 2A, 2B).

These results clearly show that D-Ala protects screens the N-terminal from degradation and could be an alternative to Aib as no degradation was detectable when D-Ala is inserted. As seen for both GLP-1 and GLP-2, DPP-IV is able to cleave with other residues (in this case serine) at the penultimate position other than the preferred proline or alanine, albeit more slowly (3-fold for GLP-1 and 57-fold for GLP-2 compared to alanine) [28]. We observed different hydrolysis rates of GLP-1 and GLP-2 despite their similar N-terminal sequence. By looking at the NMR structure of both GLP-1 (PDB 4ADP) and GLP-2 (PDB 2L63), it is evident that the N-terminal flanking region for GLP-2 is bent towards the helix, which could sterically hinder the DPP-IV enzyme and affect GLP-2’s binding to the active site. This will result in decreased proteolysis rate and thereby increased half-life. The differences in the proteolysis rates between GLP-1 and GLP-2 are also seen *in vivo* as the whole body metabolism of GLP-2 is significantly slower than for GLP-1, with half-lives of 7 min compared to 1-2 min for GLP-1 [29, 30]. Approximately after 5 min of GLP-1 proteolysis, DPP-IV starts to cleave off the two following amino acids in the sequence (EG) (data not shown).

To investigate how D-Ala would affect the potency of the corresponding GLP-1 and GLP-2 analogues, the *in vitro* receptor potency of both analogues was assessed on their respective receptors (Figure 2C, 2D). For GLP-1, a minor effect was observed when L-Ala was substituted with D-Ala, however, substituting L-Ala with L-Ser decreased the GLP-1 receptor potency around 5-fold. The opposite was observed for GLP-2, where the L-Ser substitution resulted in a more potent analogue compared to the D-Ala analogue.

The inhibition of degradation by DPP-IV and the retained potency when L-Ala is substituted with D-Ala provided us with a strong incentive to pursue GLP-1 and GLP-2 with an enzymatically inserted D-Ala.

### ProcM dehydrates serine and threonine residues in the N-terminal of GLP-1 and GLP-2

In the work by Yang and van der Donk [17], D-Ala is introduced into a core peptide by two steps, first, a LanM recognizes a specific leader peptide that guides ATP-dependent phosphorylation of serine and threonine residues, and subsequently acid-base catalyzed dehydration occurs to yield dehydroalanine and dehydrobutyrine in the following core peptide. Secondly, the Zn-dependent enzyme NpnJ hydrogenates dehydroalanine selectively and leaves the non-proteogenic D-Ala. The literature contains several other examples of using LanM enzymes on modified substrates, but our goal was to alter a specific position and leave no traces behind. This imposes several restrictions. All constructs were made in pETDuet expression vectors to enable co-expression of the peptide and the dehydratase. We needed a purification tag, the leader peptide guiding the enzymatic dehydration, a selective proteolytic site, and intact GLP-1/GLP-2 without other modifications than D-Ala. The final design and workflow can be seen in Figure 3A, 3B. We chose an N-terminal hexahistidine tag because of its size and simple use. After the purification tag the respective leader sequence for one of the six LanM enzymes (kindly provided by the van der Donk Lab) was inserted. Next to the leader sequence, we inserted a slightly altered recognition site for the tobacco etch virus protease (TEV). The most commonly used sequence is ENGLYFQ-G/S, but this would leave a glycine/serine as the N-terminal amino acid and cause either an additional Ser or Gly in the N-terminal of the final peptide or the loss of the His normally present. Neither option was acceptable, so instead, we used ENGLYFQ-H, which has been reported as an acceptable alternative [31]. Finally, we included the sequence of GLP-1/GLP-2. We used In-Fusion cloning technology to assemble the plasmids, and expression was carried out in *E.coli* BL21(DE3). The modified peptides were separated from the cell lysate using Ni-NTA pull-down, cleaved with TEV protease, and analyzed using HPLC-MS/MS. When comparing product distribution, we assumed that the modification introduced did not drastically alter the ionization properties of the peptide. We tested all six LanMs (LctM, CylM, HalM1, HalM2, NpnM, and ProcM) provided by the van der Donk lab, and perhaps unsurprisingly the degree of dehydration in GLP-1/GLP-2 varied a lot (Table S1). Because we performed everything in *E. coli,* the results are a combination of the level of expression and potential enzymatic modifications. This is clear from the LctM and HalM1 constructs, where we do not observe the precursor peptides in the *E.coli* culture and therefore can not conclude if the enzymes could accept the unnatural substrate. For NpnM, we observe expression of the precursor, but MS/MS analysis showed no sign of dehydration. For CylM and HalM2, we observed unmodified GLP-1 core peptides as well as core peptides with one and two dehydrations; however, these dehydrations are exclusively found on Thr11 and Thr13. CylM also showed signs of dehydration on Ser5 in GLP-2. Others have shown that the order and efficiency of the post-translational modifications are altered *in vitro* if the leader peptide and core peptide are added separately and hypothesized structural elements in the core peptide are important for binding the dehydratase long enough to ensure both phosphorylation and dehydration [13]. These five LanMs are known to act on one or two natural substrates, which could explain the observed modest activity on the unnatural GLP-1 and GLP-2 substrates.

**Figure 3.**
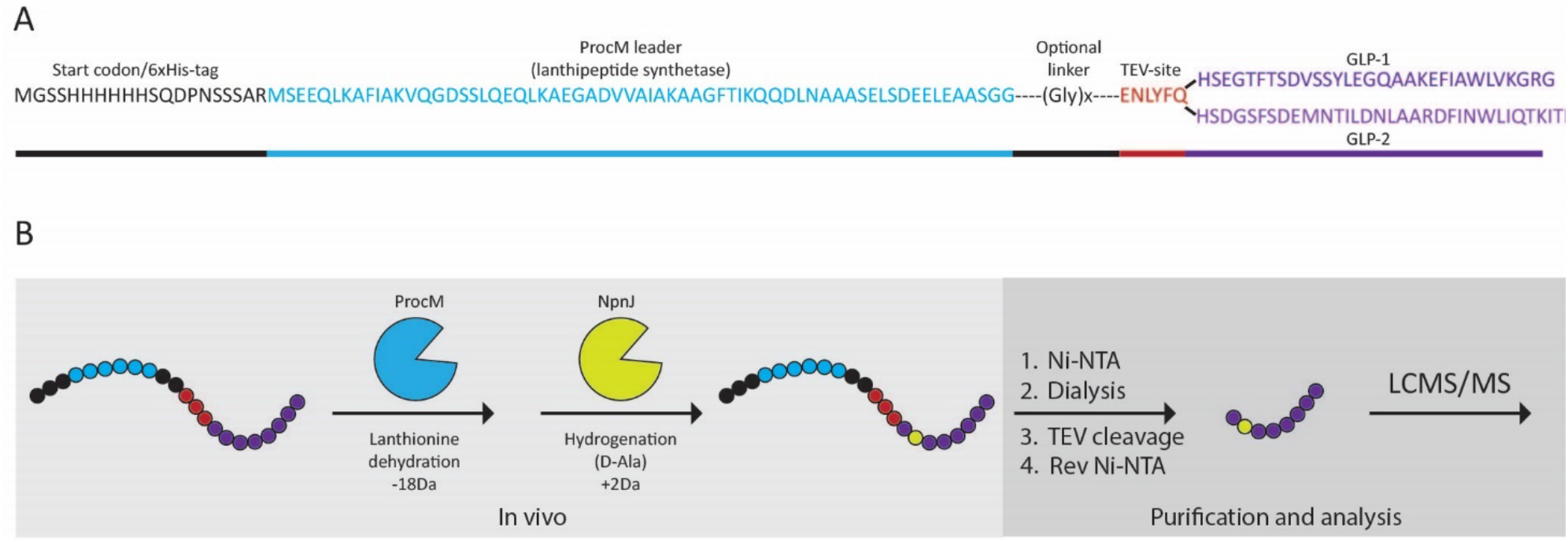
Design of constructs and workflow for peptide production and analysis. A) Sequence of GLP-1 and GLP-2 including N-terminal His-tag, leader peptide (blue), optional linker (black), TEV-site (red), and GLP-1/GLP-2 (purple). B) Workflow including co-expression of GLP-1 and GLP-2 with ProcM and NpnJ (optional) followed by purification by Ni-NTA chromatography, TEV cleavage, and reverse Ni-NTA. Purified peptides were analyzed by HPLC MS/MS for the degree of dehydration and D-Ala.

In contrast, the sixth LanM tested, ProcM, has 30 naturally occurring precursor peptides, ProcAs, with a rather conserved leader sequence [32]. With that many substrates to process ProcM must be a very promiscuous LanM enzyme [33]. We used the full leader sequence of ProcA3.3 for our expression. The van der Donk lab has previously shown that it is possible to use a truncated version of the leader in combination with another leader sequence [4]. Still, we were not able to obtain expression of GLP-1/GLP-2 with the truncated version. The promiscuity of ProcM was obvious, as we detected dehydrobutyrine and dehydroalanine residues in both GLP-1 and GLP-2 - two peptides with no sequence similarity to the natural substrates. The distribution of dehydrated residues is shown in Figure 4A, 4B. Even though the sequence of GLP-1 and GLP-2 are somewhat similar in the N-terminal, we observed a difference in the ratio of Dha and Dhb. All expressions are done in both LB media and autoinduction media, but we found that expression in autoinduction media resulted in a higher number of modifications. For every figure shown, a complementary figure can be found in the supporting information showing the outcome in LB media compared to autoinduction media. In addition, we also observed several unidentified species that could be associated with the GLP-1/GLP-2 sequence. These are shown in Supporting Figure S1-S3 together with a proposed explanation.

**Figure 4.**
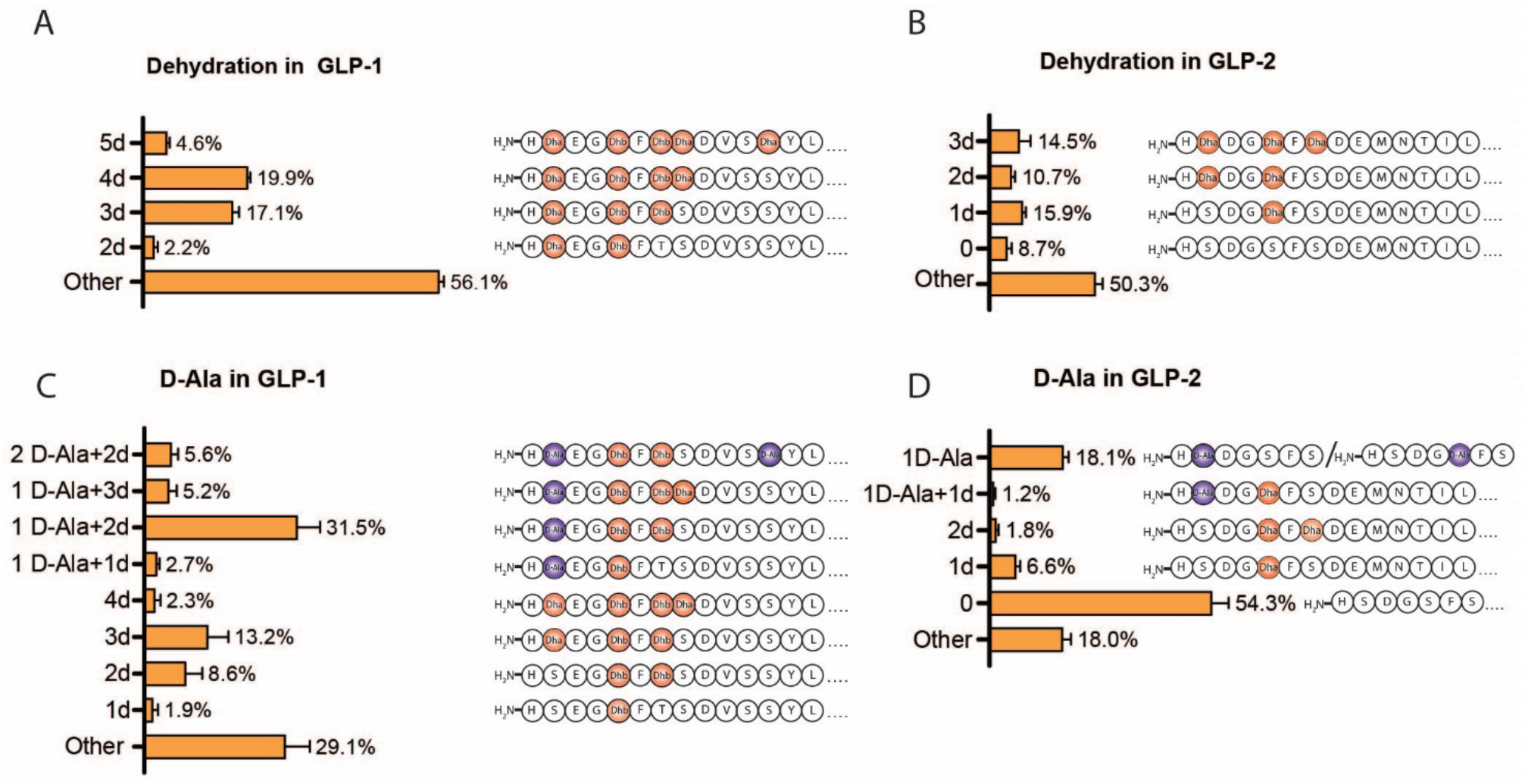
In vivo modification of GLP-1 and GLP-2. A+B) Bar chart (left) showing % distribution of dehydrated GLP-1 and GLP-2 variants analysed by HPLC-MS/MS. Sequence (right) shows the position of modfied residue(s). C+D) Bar chart (left) showing % distribution of D-Ala variants of GLP-1 and GLP-2 analysed by HPLC-MS/MS. Sequence (right) shows position of modfied residue(s).

ProcM modifies all the expressed GLP-1 and almost 20% of the identified GLP-1 has incorporated four dehydrations and the remaining with two, three, and five dehydrations. Ser8 is dehydrated in all GLP-1s compared to GLP-2, where only 25% has a dehydrated Ser2, and 8.7% are not modified by ProcM. The N-terminal of the two are almost identical except for the third amino acids in each sequence (Glu9 GLP-1 and Asp3 GLP-2), but looking at the natural substrates for ProcM, Ser/Thr-residue are both found to have Glu and Asp on their C-terminal [33] and according to studies by Rink et al. there should not be a difference in the dehydration pattern with a Glu or Asp C-terminally located to the Ser [6]. It is more likely caused by small structural differences.

### Co-expression with NpnJ reduce Dha to D-Ala

To reduce the dehydroalanine to D-Ala instead of Ser8 (Ser2 in GLP-2), we co-expressed NpnJ together with ProcM and GLP-1/GLP-2. Both peptides showed some degree of reduction (Figure 4C, 4D), as 19.3% of GLP-2 had a D-Ala incorporated in the sequence. Approximately half of the incorporated D-Ala in GLP-2 was incorporated on Ser5 instead of Ser2 and 54.3% of GLP-2 were not modified in any way. Co-expressing with NpnJ affects the level of modifications observed in GLP-2 as only 8.7% were unmodified in absence of NpnJ. The additional burden caused by the introduction of NpnJ on pCDFDuet-1 could lead to lower levels of ProcM and hence fewer modifications on the core peptide, but we did not investigate this.

Dha8 in GLP-1 was reduced in 45% of the observed peptides. In order to increase the amount of D-Ala8 (GLP-1) and D-Ala2 (GLP-2), we tried to support NpnJ with chaperones (GroES and GroEL). Both chaperones have to be present when NpnJ is expressed and purified [17]. GroES and GroEL were both encoded in the same plasmid as NpnJ and co-expressed with ProcM and GLP-1/GLP-2. Co-expressing NpnJ with chaperones did not significantly increase the number of peptides with Dha8/2 in GLP-1/GLP-2 (Supporting Figure S4). Based on this preliminary data, we noticed a distance-dependent trend. For GLP-1, Ser8 would often be dehydrated, while Ser17 and Ser18 were hardly ever dehydrated. We reasoned that the distance from the leader sequence could be exploited to manipulate selectivity and designed a series of constructs with variable Gly-spacers to test this hypothesis by pushing the GLP core peptide further away from the leader sequence.

### The use of Gly-linkers changes the modification pattern

We set out to investigate the distance-dependency from the leader sequence to the Ser and Thr residues as literature for other LanMs indicates that Ser/Thr have to be placed at a certain distance downstream of the leader sequence, presumably for those residues to reach the active site of the enzyme. Chatterjee *et al.*showed that Ser4 is not modified in wild type lacticin, because it is too close to the leader peptide and flanked by two Gly residues [34]. They were able to obtain dehydration of that Ser when three Ala residues were inserted between the leader and core peptide together with mutation of the two surrounding Gly residues [34]. Similarly, others have hypothesized that the sequential modification steps can be controlled by substrate contractions caused by cyclizations, which would bring otherwise too distant Thr/Ser within reach of the dehydratase domain [35]. By looking at the crystal structure of CylM (PDB 5DZT), it is clear that there is a certain distance from the leader binding region to the active site of the dehydratase domain [36]. Taking these previous studies into consideration, we designed GLP-1/GLP-2 with a 5x and 10x Gly spacer after the leader sequence (Figure 3A). The dehydration pattern is shown in Figure 5A for GLP-1, and showed that inserting a linker prevents dehydration of Ser18. For GLP-2 (Figure 5B) the 5x Gly linker resulted in 13.8% with only one dehydration (Dha2) as the only modification. Increasing the linker further with a 10x Gly linker, the fourth dehydration is lost in GLP-1 (Figure 5C) supporting the fact that the distance from the leader peptide to the desired Ser/Thr residue is important. Increasing the linker length for GLP-2 decreased the amount of GLP-2 with a single dehydration, but counterintuitively two and three dehydrations were observed (Figure 5D). Co-expressing GLP-2 5x Gly with NpnJ did not result in any modifications (Figure 5F), in contrast to the 10x Gly linker (Figure 5H), where 18.2% of GLP-2 has a D-Ala incorporated, but it is a mix of D-Ala2 and D-Ala5. Co-expressing GLP-1 5x Gly with NpnJ did not increase the overall level of incorporated D-Ala8, but we are now able to get 3.4% of GLP-1 with D-Ala8 as the only modification in the sequence (Figure 5E) and increasing the linker further with 10x Gly, we were able to get as much as 18.8% (Figure 5G) (Supporting Figure S2+3).

**Figure 5.**
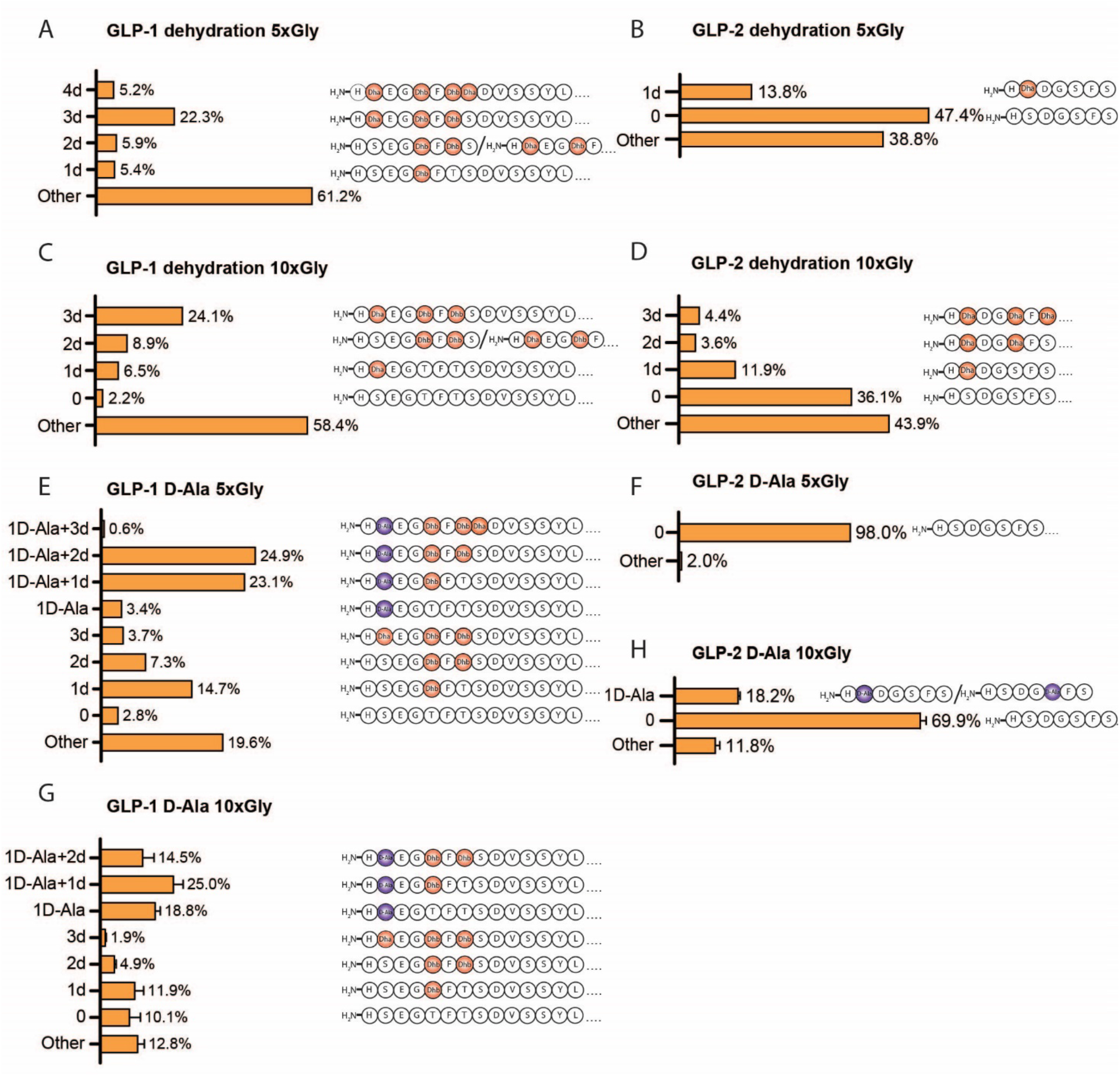
Effect of linker lenghts on in vivo modification of GLP-1 and GLP-2. Bar chart (left) showing % distribution of variants and sequence (right) showing position of modfied residues. A+B) Dehydration of GLP-1 and GLP-2 expressed with 5x Gly linker, respectively. C+D) Dehydration of GLP-1 and GLP-2 expressed with 10x Gly linker, respectively. E+F) D-Ala content of GLP-1 and GLP-2 expressed with 5x Gly linker, respectively. G+H) D-Ala content of GLP-1 and GLP-2 expressed with 10x Gly linker, respectively.

The co-expression with 5x and 10x Gly linkers was also expressed with chaperons (GroES/GroEL), but that did not have a positive effect on the D-Ala incorporation (Supporting Figure S5). We tried to increase the linker length further to 15x and 20x Gly linker (Supporting Figure S6). For GLP-2 a maximum was reached, as no modifications were observed for both linkers. For GLP-1 with 15x Gly linker the overall level of D-Ala8 decreased a bit to 51.5%, but the level of GLP-1 with D-Ala8 as the only modification decreased a lot. For the 20x Gly linker a maximum was reached as no modifications were observed. Our results clearly demonstrate that modifications are dependent on the linker length between the leader and core peptides, but also that is not entirely predictable.

## Conclusion

The introduction of post-translational modifications in therapeutic peptides could serve as an important tool to complement synthetic chemistry and protection against proteolytic clevage. The enzymes found in RiPP biosynthesis are known to introduce a large number of attractive modifications such as thioether bridges, C-terminal amides, backbone amide methylations, and the introductions of D-amino acids. In this work, we investigate how D-Ala can be introduced into a strategically important position in the hormone GLP-1 by screening a small set of LanM enzymes and their accompanying leader peptides. Importantly, this provides a direct comparison of several LanMs on the same substrates and demonstrates how glycine linkers can be used to alter the modification patterns. We show that CylM and HalM2 produced a narrow product distribution, but with a preference for Thr11 instead of the desired Ser8 in GLP-1. We show that linkers inserted between the leader and the designed core peptide can be utilized to modulate product distribution. Finally, we achieve a substantial fraction of the desired product by using the promiscuous ProcM. However, the broad distribution of different products with nearly identical retention times on RP-HPLC represents a considerable challenge that needs further attention to be solved.

## Methods

### Cloning

Plasmids used in this study were constructed using In-Fusion^®^ Cloning Technology (TaKaRa). Primers and genes were ordered from Integrated DNA Technologies (IDT). Linear plasmid fragments were generated by PCR and purified by Monarch^®^ PCR and DNA cleanup kit (New England BioLabs^®^). Ligations were performed with In-Fusion^®^ enzyme at 50°C for 15 minutes (min) (50 ng vector and 3-fold excess of insert). *E. Coli* DH5α chemically competent cells were transformed with the generated plasmids. Positive clones were selected for on LB agar with the appropriate antibiotic. All sequences were validated by Sanger sequencing. All plasmids used in this study are listed in Table S2 and were constructed using primers listed in Table S3.

Plasmids coding for ProcM[4], NpnM[32], LctM[37], CylM[38], HalM1[39], HalM2[39] and NpnJ[17] were kindly provided by the van der Donk lab (Table S2). ProcM was received in pACYCDuet-1 with an empty MCSI, this was not further engineered. NpnJ was received fused to 6x His-tag, MBP and a thrombin site in pRSFDuet-1, this was not further modified. NpnJ was also received in pCDFDuet-1 without a 6x His-tag, here we inserted the two chaperones GroES and GroEL from the pGro7 plasmid (TaKaRa). The sequence for GroES and GroEL were amplified from the pGro7 plasmid by PCR and inserted into the linearized plasmid with NpnJ by In-Fusion^®^. LctM, NpnM, HalM1, HalM2 and CylM were all amplified by PCR and inserted into an empty pACYCDuet-1 vector.

To insert the respective leader matching each enzyme, plasmids were cleaved with FD *SgsI* (Thermo Scientific^™^) and treated with FastAP termosensitive alkaline phosphatase (Thermo Scientific^™^). DNA encoding leader sequences were ordered as gene blocks and ligated into the plasmids using In-fusion^®^. DNA encoding enzymes were placed in MCSII in the plasmids, and DNA encoding 6x His-tag, leader sequence, TEV site and GLP-1/GLP-2 were placed in MCSI. Plasmids encoding enzyme and the respective leader sequence were linearized by PCR and gene blocks of GLP-1 and GLP-2 were inserted using In-fusion^®^.

### Cultivation and expression of GLP-1 and GLP-2

Chemically competent *E. coli* BL21(DE3) cells were transformed with the different constructs. A single colony was used to inoculate LB medium (10-20 mL) supplemented with either kanamycin (50 μg/mL), chloramphenicol (34 μg/mL), or streptomycin (50 μg/mL) (or both chloramphenicol and streptomycin when co-expressing two plasmids) and then cultivated overnight. The starter culture was used to inoculate a flask containing LB/Autoinduction media (Table S6) (100 mL). Cells were grown to an OD600 of ~ 1-1.5 (37°C), and the expression in LB media was induced with 0.5mM IPTG overnight and transferred to 18°C together with the expression in Autoinduction media. Cells were harvested by centrifugation (6000 rpm, 15 min, 4°C). The pellet was resuspended, lysed in B-Per^™^ (Thermo Scientific^™^), and incubated at 4°C (10 min). Cell debris was pelleted via centrifugation (12000 rpm, 30 min, 4°C).

### Protein purification

The supernatant was incubated with Ni-NTA beads (30 min, 4°C) and afterwards applied to Micro BioSpin^™^ Columns (0.8mL). Beads were washed with three times the volume of the beads with wash buffer (50 mM HEPES, 500 mM NaCl, 10 mM Imidazole, pH 7.5). Proteins were eluted with elution buffer (50 mM HEPES, 500 mM NaCl, 400 mM Imidazole, pH 7.5). The eluted proteins were visualized on SDS-PAGE and the elution were dialysed against PBS overnight (4°C). TEV was added in 1:10 w/w ratio and incubated overnight (4°C). The cleaved proteins were incubated with Ni-NTA beads for revers his-purification and cleaved proteins were collected in the flow through and visualized on SDS-PAGE (Mini Protean Tris-Tricine Precast gels, BioRad).

### Peptide synthesis and purification

All peptides were prepared by solid phase peptide synthesis using the Fmoc strategy and dissolved in dimethylformamide (DMF)+4% oxyma. Rink-amide or preloaded Wang resin (both at 0.1 mmol) was used and dissolved in 10 mL DMF. All reactions were performed using the Liberty blue HT12 (CEM). After synthesis, the resin was washed with dichloromethane (DCM) (6×10 mL) to remove remaining DMF. To release the peptide from the resin and to remove all protecting groups on the side chains, Triisopropylsilane (TIPS) (25 mL 2.5%), DTT (2.5% w/v) and H2O (2.5%) in trifluoracetic acid (TFA) was mixed with the resin and incubated while shaking (2 hours). The cleaved peptide was precipitated using ice-cold Diethyl ether (30 mL) and pelleted by ether centrifugation (3500 rpm, 3 minutes). The pellet was washed twice in diethyl ether and stored at −20 °C until purification. A pinch of each pellet was dissolved in 1:1 acetonitrile (MeCN) and H2O (300 μL) and used for LC-MS (mass accuracy) and UHPLC (purity) analysis. LC-MS was performed using a C18 BEH column (Waters) with a 5-95% MeCN gradient. Masses was detected using a Xevo G2-XS QTof (Waters). UHPLC was also performed using a C18 BEH column and eluted with a gradient of MeCN+0.05% TFA.

Precipitated peptides were resuspended in Acetic acid diluted in MilliQ H_2_O (40%) and purified by RP-HPLC on a C18-column equilibrated in buffer A (0.1% TFA in H_2_O) with 10% buffer B (0.1% TFA in MeCN). After sample application, the column was washed with buffer A (250 mL) before a gradient of B (0.1% TFA in MeCN) was used for elution. Gradients was optimized for each peptide for better separation. Fractions were collected based on UV at 230 nm. The purity of each fraction was analysed by UHPLC and the best fractions were pooled and tested again on LC-MS and UHPLC to get the final purity. Some samples were not pure enough after RP-HPLC, these were purified again at neutral pH. The sample was applied to a column equilibrated in buffer C (1% (NH_4_)_2_CO_3_) and eluted with a gradient of buffer D (100% MeCN). Fractions were analyzed by UHPLC, the best fractions were pooled and purified again using buffer A and B as described. The best fractions were pooled and analyzed and the final concentration was measured by charged aerosol detection (CAD). Aliquots were lyophilized and stored at −80 °C.

### MS/MS characterization

RP-HPLC coupled with tandem mass spectrometry (RP-HPLC-MS/MS) was performed using a Waters Synapt G2S-i instrument (Wilmslow, UK), equipped with a Waters Acquity UPLC CSH C18 (1.7μm, 130Å, 1.0×150 mm), water (eluent A), acetonitrile (eluent B) (both containing 0.1% formic acid and premixed from Fisher Chemicals, Thermo). Mass spectrometry was done in positive electrospray mode (+3kV) in high resolution mode with a nominal resolution of 30.000. For fragmentation, expressed mass spectrometry (MSE) was done with a collision energy ramp of 25-45V. Data analysis was conducted using MassLynx 4.2. Masses were extracted from all the peaks observed in the Total Ion Chromatogram (TIC), the most abundant m/z value for each peptide of interest was extracted (XIC) and integrated. For analyzing fragmentation data, the mass spectra of interest was deconvoluted using MaxEnt 3 and observed masses was compared with theoretical masses.

### Transfection for cAMP potency assay for GLP-2

One day prior to the transfection, HEK293T cells were seeded in a T25 cell culture flask (1.200.000 cells/flask) and cultured in 5 ml DMEM (Dulbecco’s modified Eagle’s medium) 1885 medium, supplemented with 10% fetal bovine serum (FBS), and 1% penicillin (180 U/ml)/streptomycin (45 μg/ml) and incubated at 37°C, 5% CO_2_, and 95% air humidity. The cells were transiently transfected using the calcium phosphate precipitation method. 10 μg of WT GLP-2R was added to a total volume of 120 μl in Tris-EDTA (TE) buffer (10 mM Tris-HCl, 2 mM EDTA-Na_2_, pH 7.5). Thereafter, 15 μl CaCl_2_ (2 M) was added to the mix before adding the mixture dropwise into 120 μl 2 x Hepes-buffer saline (HBS) buffer (280 mM NaCl, 50 mM Hepes, 1.5 mM Na_2_HPO_4_, pH 7.2). The final transfection mix was incubated for 45 min before adding dropwise to the cells. The transfection was terminated after 5 hours by replacing the transfection medium with 5 ml supplemented DMEM.

### GLP-2R signaling assay, cAMP measurements

One day prior to the assay (Discover HitHunter cAMP assay), the transfected cells were seeded in a white 96-well cell culture microplate (25.000 cells/well). The assay was performed by washing the cells with 100 μl 1xHBS and subsequently incubating the cells for 30 min in 100 μl IBMX (3-isobutyl-1-methylxanthine) diluted in 2xHBS at 37°C. Thereafter, 5 μl ligand in concentrations ranging from (1 pM to 1 μM) was added to the plate and incubated for 30 min at 37°C. After incubation, the ligand and IBMX were aspirated, and 30 μl PBS, 10 μl cAMP antibodies, and 40 μl (enzyme donor/lysis buffer/CL) were added and incubated for 60 min at room temperature in the dark before adding 40 μl enzyme acceptor solution. After 3 hours incubation at room temperature in the dark, the cAMP was measured as luminescence using PerkinElmer EnVision 2104 Multilabel Reader. EC_50_-values were calculated by non-linear curve fitting using a three parameter logistic model with shared basal response using GraphPad Prism (version 9.0.1). The results were normalized to the efficacy of the native GLP-2 hormone.

### GLP-1R signaling assay, cAMP potency

GLP-1R potency (cAMP) was measured using a clonal BHK cell line co-expressing the human GLP-1R and the cAMP response element (CRE) luciferase reporter gene. On the day of the assay, frozen cells stocks of assay-ready cells were thawed in a 37°C water bath, washed once in PBS (Gibco, 14190-094) and diluted to 100.000 cells/mL in assay buffer consisting of DMEM without phenol red (Gibco, 11880-028) supplemented with 1X GlutaMAX (Gibco, 35050-038), 10 mM HEPES (Gibco, 15630-056), 1% (w/v) ovalbumin (Sigma, A5503) and 0.1% (v/v) Pluronic F-68 (Gibco, 24040-032). Serial dilutions (10-fold dilutions, 8 concentrations pr. compound) were performed in assay buffer without in a 96-well dilution plate (Greiner Bio-one, U-shape, 650201) using a Biomek i7 liquid handler (Beckman-Coulter). From the dilution plate, 50 μL of each dilution was transferred to 96-well assay plates (ThermoFisher, 237105) to which 50 μL of the cell suspension was added (5.000 cells/well). The assay plates were incubated for 3 hours at 37°C in 5% CO_2_, left at room temperature for 5 minutes after which 100 μL SteadyLite Plus (PerkinElmer, 6066759) was added to each well. Plates were sealed and incubated at room temperature with gentle shaking for 30 minutes while protected from light. Luminescence was detected on a Synergy 2 (BioTek) luminescence plate reader and the EC_50_-values calculated by non-linear curve fitting using a three parameter logistic model with shared basal response using GraphPad Prism (version 9.0.1). The results were normalized to the efficacy of the native GLP-1 hormone.

### DPP-IV proteolysis assay

Synthesized peptides were diluted in assay buffer (0.005% Tween20 in PBS buffer pH 7.2) (250 μM). The reaction was carried out in assay buffer, 10 nM DPP-IV and 25 μM peptide. Samples were taken at 0, 5, 15, 30, 60, 120, 240, 360, 960, 1440 min and the reaction was stopped by mixing incubate (30 μL) with methanol (99%)+1% formic acid (90 μL) and samples were analysed on RP-HPLC MS/MS.

## Acknowledgement

This work was financially supported by an Industrial Ph.D. Project Grant from Innovation Fund Denmark (8053-00220B).

