## Supporting Information for "Using LanM Enzymes to Modify Glucagon-Like Peptides 1 and 2 in *E.coli*"

### Supplementary

| | **GLP-1** | **GLP-2** |
|---|---|---|
| **LctM** | * | * |
| **NpnM** | 0 dehdyrations | 0 dehydrations |
| **CylM** | 0 dehydrations<br>1 dehydration (Thr11)<br>2 dehydrations (Thr11+Thr13) | 0 dehydrations<br>1 dehydration (Ser5) |
| **HalM1** | * | * |
| **HalM2** | 0 dehydrations<br>1 dehydration (Thr11)<br>2 dehydrations (Thr11+Thr13) | 0 dehydrations |

*Table S1 – Five of the provided enzymes and their dehydration pattern in GLP-1 and GLP-2. Boxes marked with "*" indicate that GLP-1/2 were not expressed.*

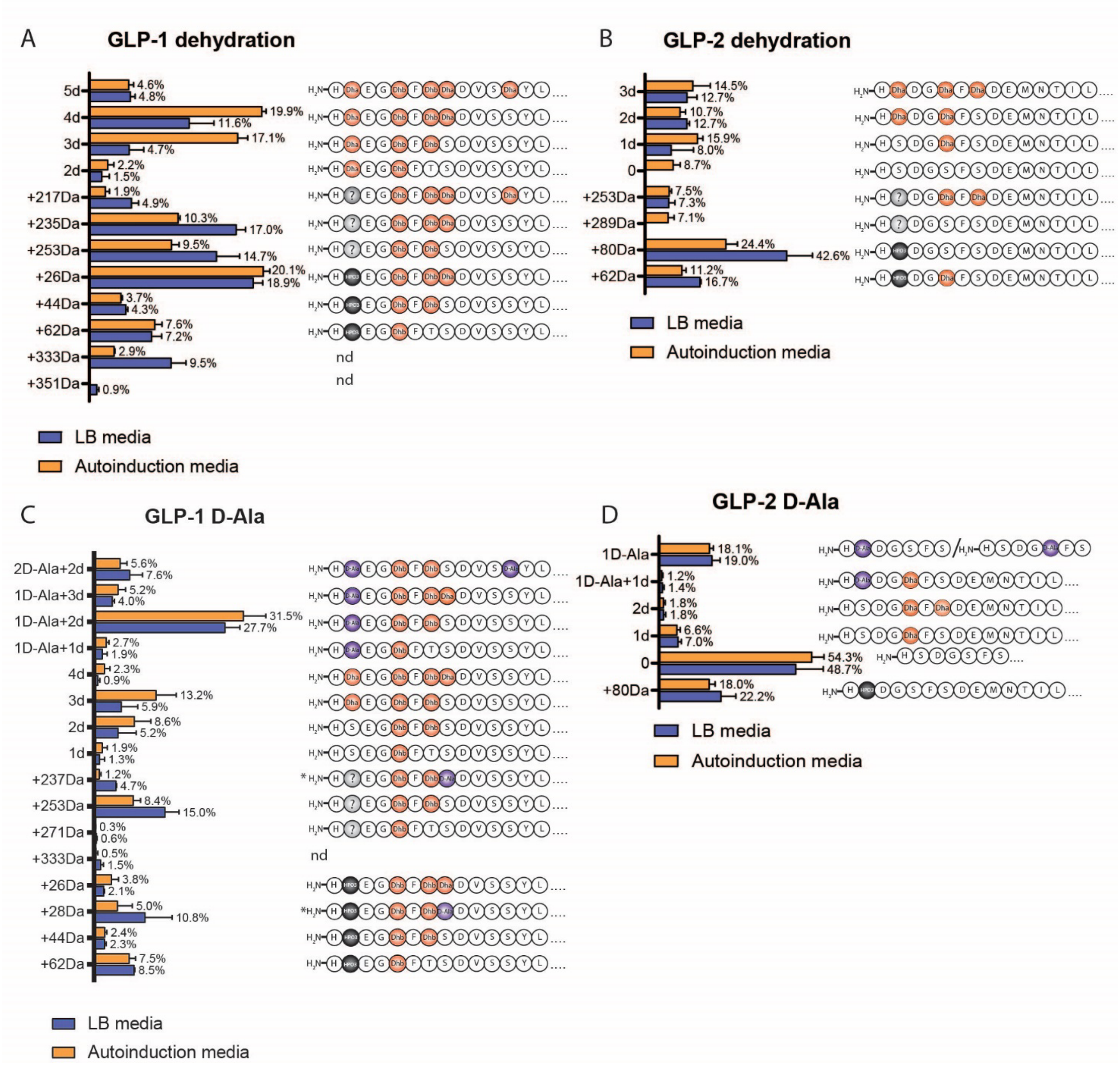


*Figure S1 – In vivo modification of GLP-1 and GLP-2 expressed in LB and autoinduction media, all done in triplicates. A+B) Bar chart (left) showing % distribution of dehydrated GLP-1 and GLP-2 variants analyzed by HPLC-MS/MS. Undefined species marked as "other" in the main text are shown and the modified sequence is shown to the right. C+D) Bar chart (left) showing % distribution of D-Ala variants of GLP-1 and GLP-2 analyzed by HPLC-MS/MS and sequences with modifications are shown to the right.*

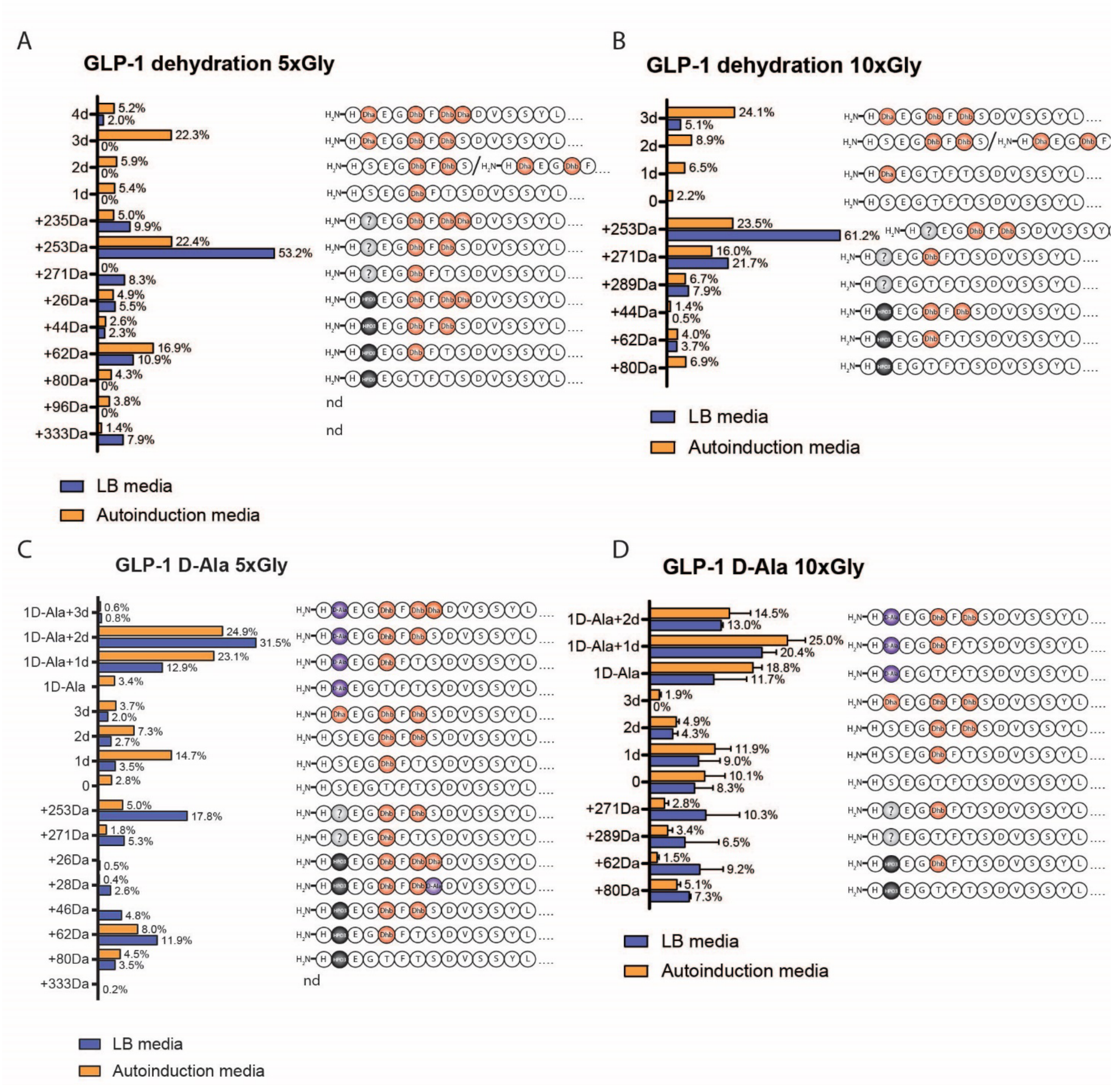


*Figure S2 – In vivo modification of GLP-1 expressed in LB and autoinduction media with 5x and 10x Gly-linker. Bar chart (left) showing % distribution of variants and the modified sequences are shown to the right. A) Distribution of dehydration in GLP-1 expressed with a 5x Gly linker. B) Distribution of dehydration in GLP-1 expressed with a 10x Gly linker. C) Distribution of dehydration and D-Ala in GLP-1 expressed with a 5x Gly linker. D) Distribution of dehydration and D-Ala in GLP-1 expressed with a 10x Gly linker, done in triplicate.*

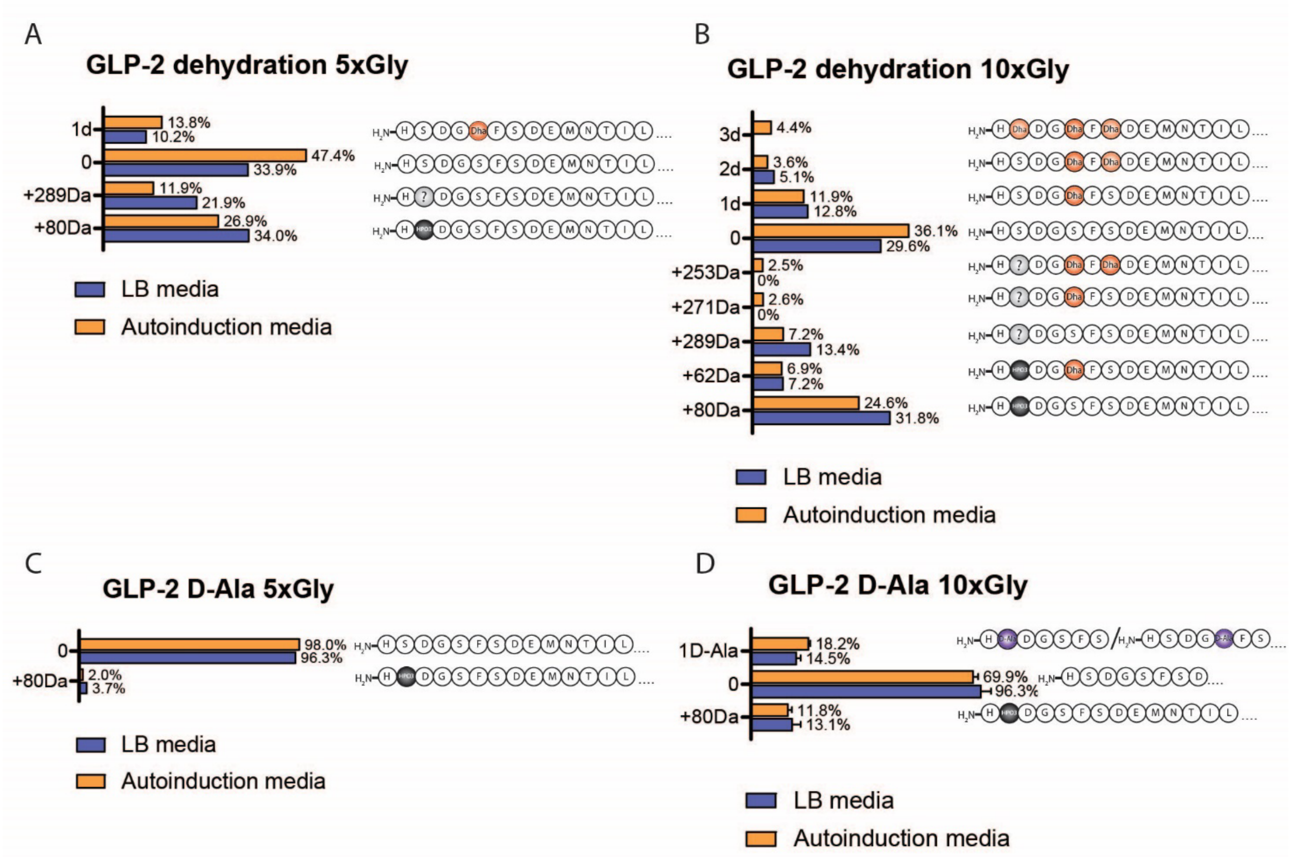


*Figure S3 – In vivo modification of GLP-2 expressed in LB and autoinduction media with 5x and 10x Gly-linker. Bar chart (left) showing % distribution of variants and the modified sequences are shown to the right. A) Distribution of dehydration in GLP-2 expressed with a 5x Gly linker. B) Distribution of dehydration in GLP-2 expressed with a 10x Gly linker. C) Distribution of dehydration and D-Ala in GLP-2 expressed with a 5x Gly linker. D) Distribution of dehydration and D-Ala in GLP-2 expressed with a 10x Gly linker, done in triplicate.*

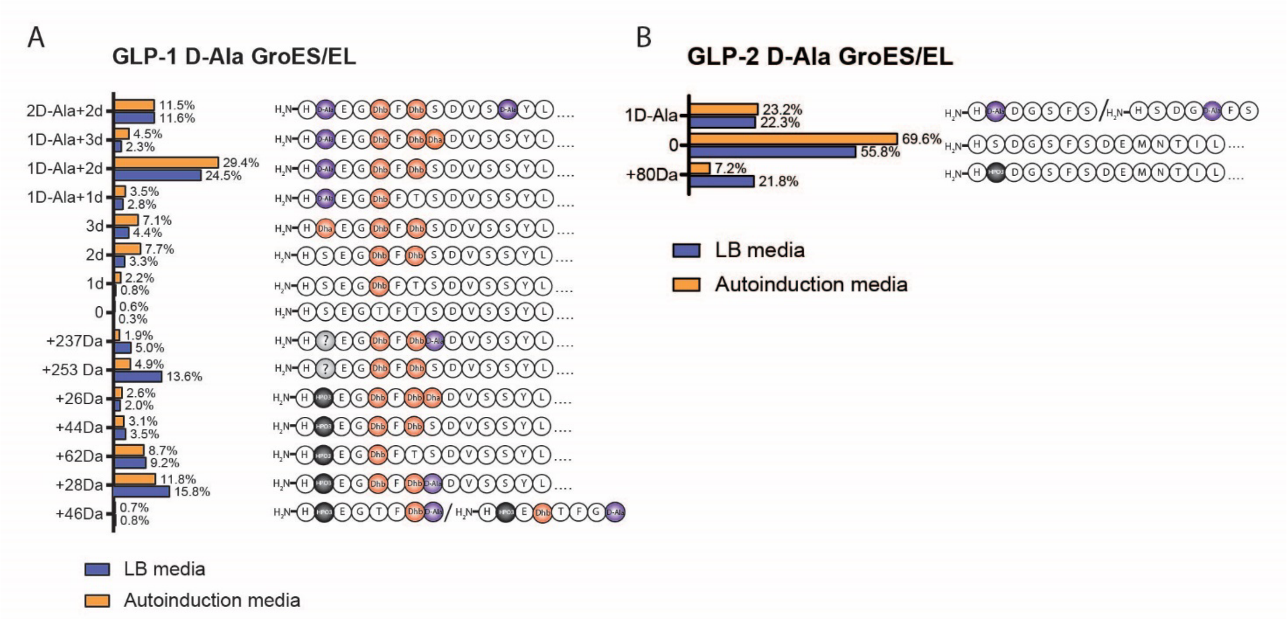


*Figure S4 – In vivo modification of GLP-1 and GLP-2 expressed in LB and autoinduction media with chaperons (GroES/EL). Bar chart (left) showing % distribution of variants and the modified sequences are shown to the right A) GLP-1. B) GLP-2*

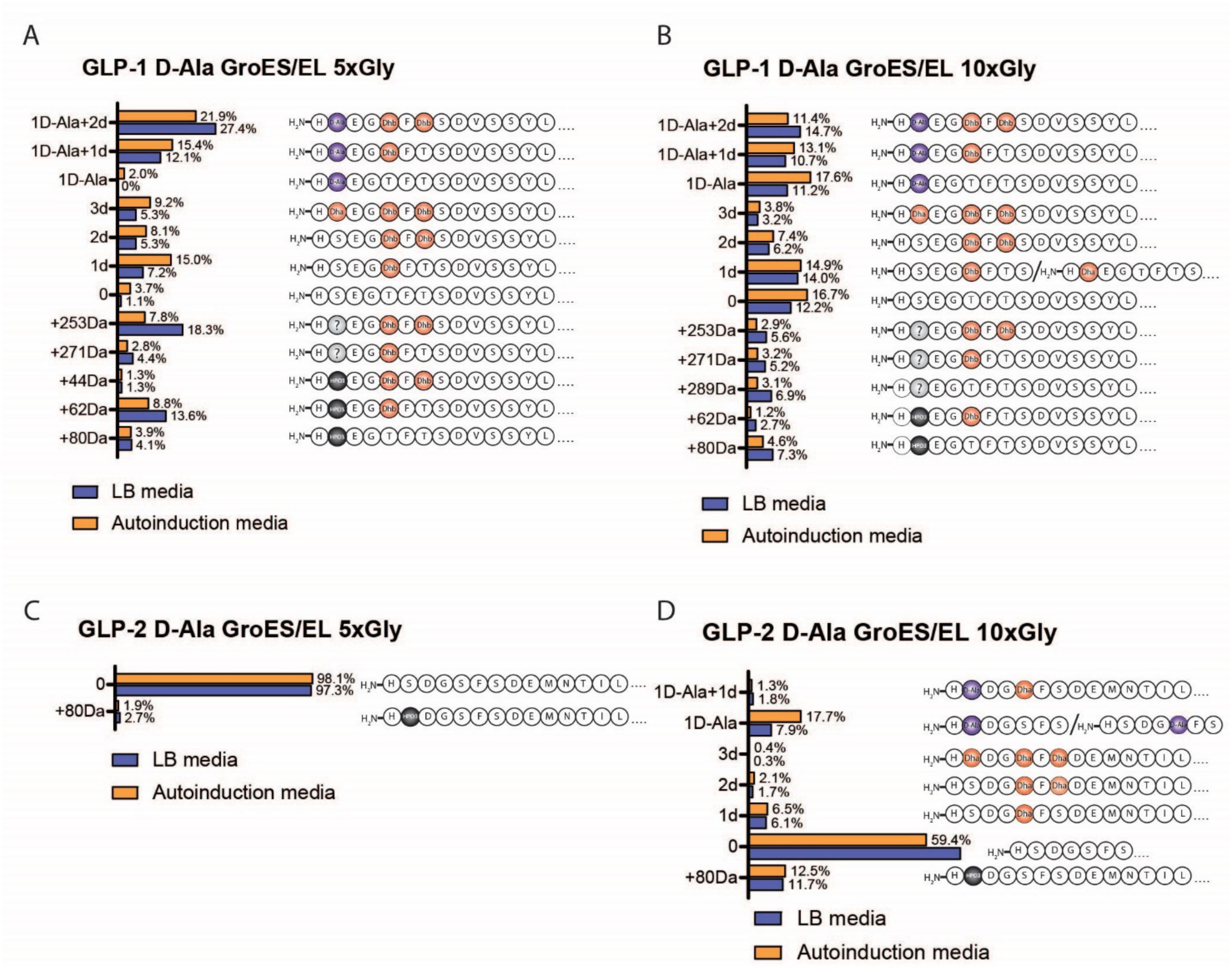


*Figure S5 – In vivo modification of GLP-1 and GLP-2 expressed in LB and autoinduction media with 5x and 10x Gly linkers and with chaperons (GroES/EL). Bar chart (left) showing % distribution of variants and the modified sequences are shown to the right A) GLP-1 5x Gly linker. B) GLP-1 10x Gly-linker. C) GLP-2 5x Gly linker D) GLP-2 10x Gly linker.*

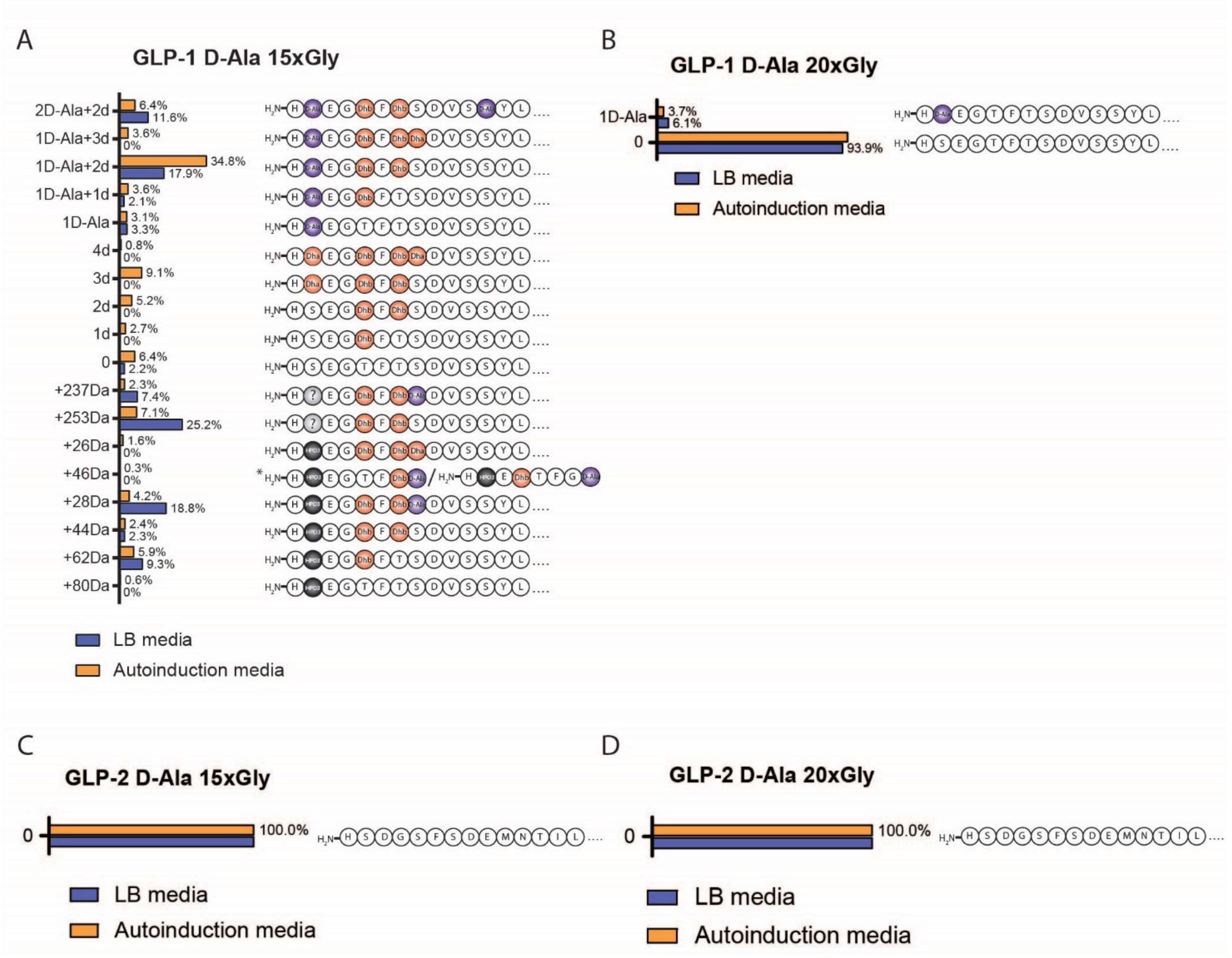


*Figure S6 – In vivo modification of GLP-1 and GLP-2 expressed in LB and autoinduction media with 15x and 20x Gly-linker. Bar chart (left) showing % distribution of variants and the modified sequences are shown to the right. A) GLP-1 15x Gly linker. B) GLP-1 20x Gly linker. C) GLP-2 15x Gly linker. D) GLP-2 20x Gly linker.*

| No. | Name | Plasmid backbone | Genes<br>MCSI : MCSII | |
|---|---|---|---|---|
| 1 | ProcM | pACYCDuet-1 | Empty : ProcM | [1] |
| 2 | LctM | pET28b | - | [2] |
| 3 | CylM | pRSFDuet-1 | - | [3] |
| 4 | HalM1 | pET28b | - | [4] |
| 5 | HalM2 | pET28b | - | [4] |
| 6 | NpnM | pRSFDuet-1 | NpnA3 : NpnM | [5] |

| | | | | |
|---|---|---|---|---|
| 7 | NpnJ1 | pRSFDuet-1 | His6-MBP-NpnJ : empty | [6] |
| 8 | NpnJ2 | pCDFDuet-1 | MBP-NpnJ : empty | [6] |
| 9 | pACYC-1-GLP1 | pACYCDuet-1 | GLP-1 : ProcM | This study |
| 10 | pACYC-1-GLP2 | pACYCDuet-1 | GLP-2 : ProcM | This study |
| 11 | pCDF-chap | PCDFDuet-1 | MBP-NpnJ : GroES/GroEL | This study |
| 12 | pCDF-NpnJ | pCDFDuet-1 | MBP-NpnJ : empty | This study |
| 13 | pACYC-5x-GLP1 | pACYCDuet-1 | 5xGly-GLP-1 : ProcM | This study |
| 14 | pACYC-5x-GLP2 | pACYCDuet-1 | 5xGly-GLP-2 : ProcM | This study |
| 15 | pACYC-10x-GLP1 | pACYCDuet-1 | 10xGly-GLP-1 : ProcM | This study |
| 16 | pACYC-10x-GLP1 | pACYCDuet-1 | 10xGly-GLP-2 : ProcM | This study |
| 17 | pACYC-15x-GLP1 | pACYCDuet-1 | 15xGly-GLP-1 : ProcM | This study |
| 18 | pACYC-15x-GLP2 | pACYCDuet-1 | 15xGly-GLP-2 : ProcM | This study |
| 19 | pACYC-20x-GLP1 | pACYCDuet-1 | 20xGly-GLP-1 : ProcM | This study |
| 20 | pACYC-20x-GLP1 | pACYCDuet-1 | 20xGly-GLP-2 : ProcM | This study |
| 21 | pACYC-1 | pACYCDuet-1 | Empty : ProcM | This study |
| 22 | pACYC-leader | pACYCDuet-1 | 6xhis-leader-TEV : ProcM | This study |
| 23 | pRSF-NpnM-1 | pRSFDuet-1 | GLP-1 : NpnM | This study |
| 24 | pRSF-NpnM-2 | pRSFDuet-1 | GLP-2 : NpnM | This study |
| 25 | pACYC-CylM-1 | pACYCDuet-1 | GLP-1 : CylM | This study |
| 26 | pACYC-CylM-2 | pACYCDuet-1 | GLP-2 : CylM | This study |
| 27 | pACYC-HalM1-1 | pACYCDuet-1 | GLP-1 : HalM1 | This study |
| 28 | pACYC-HalM1-2 | pACYCDuet-1 | GLP-2 : HalM1 | This study |

| | | | | |
|---|---|---|---|---|
| 29 | pACYC-HalM2-1 | pACYCDuet-1 | GLP-1 : HalM2 | This study |
| 30 | pACYC-HalM2-2 | pACYCDuet-1 | GLP-2 : HalM2 | This study |
| 31 | pACYC-LctM-1 | pACYCDuet-1 | GLP-1 : LctM | This study |
| 32 | pACYC-LctM-2 | pACYCDuet-1 | GLP-2 : LctM | This study |

*Table S2 - **Plasmids used for this study**. The design of plasmids either used alone or in combination. Plasmid 9,10,13-20 were expressed alone and in combination with 7, 8 or 11. Plasmid 13-20 were only expressed in combination with 8. Plasmid 9,10,13-20,22-32 had 6x his-tag, a respective leader, and a TEV in front of the GLP-1/2 sequence. Abbreviations: No, number; MCS, multiple cloning site; MBP, maltose-binding protein.*

| **Primer name** | **Primer sequence (5' to 3')** | **Purpose** |
|---|---|---|
| GroES/GroEL F | agatctcaattggatatgaatattcgtccattgcatgatc | Amplified from pGro7 w. overhangs to pCDF-NpnJ |
| GroES/GroEL R | gcgtggccggccgatttacatcatgccgcccat | Amplified from pGro7 w. overhangs to pCDF-NpnJ |
| NpnJ F | atcggccggccacgcgat | Linearize pCDF-NpnJ to insert GroES/GroEL by In-Fusion |
| NpnJ R | atccaattgagatctgccatatgtatatctcc | Linearize pCDF-NpnJ to insert GroES/GroEL by In-Fusion |
| pACYC-leader F | tagcgcgcctgcaggtcg | Linearize pACYC-leader to insert GLP-1/2 by In-Fusion |
| pACYC-leader R | ttggaagtacaggttctcaccccc | Linearize pACYC-leader to insert GLP-1/2 by In-Fusion |
| Linker-1 F | gagaacctgtacttccaacact | Linearize pACYC-1-GLP1 to insert linkers |
| Linker-2 F | gagaacctgtacttccaacattct | Linearize pACYC-1-GLP2 to insert linkers |
| Linker R | accccccgaagcagcttc | Linearize pACYC-1-GLP1/GLP-2 to insert linkers |
| 5xGly F | gctgcttcggggggtggtggcggaggaggcgagaacctgtacttc | Used as insert into pACYC-1-GLP-1/2 by In-Fusion |
| 5xGly R | gaagtacaggttctcgcctcctccgccaccaccccccgaagcagc | Used as insert into pACYC-1-GLP-1/2 by In-Fusion |
| 10xGly F | gctgcttcggggggtggtggaggcgggggaggcggcggtggtgg agagaacctgtacttc | Used as insert into pACYC-1-GLP-1/2 by In-Fusion |
| 10xGly R | gaagtacaggttctctccaccaccgccgcctcccccgcctccacca ccccccgaagcagc | Used as insert into pACYC-1-GLP-1/2 by In-Fusion |
| 15xGly F | gctgcttcggggggtggtggcggtggcggtggtggtggaggaggc ggaggcgggggaggcgagaacctgtacttc | Used as insert into pACYC-1-GLP-1/2 by In-Fusion |

| | | |
|---|---|---|
| 15xGly R | gaagtacaggttctcgcctcccccgcctccgcctcctccaccaccac cgccaccgccaccaccccccgaagcagc | Used as insert into pACYC-1-GLP-1/2 by In-Fusion |
| 20xGly F | gctgcttcggggggtgggggcggtggagggggtggagggggagg cggaggcggcgggggtggaggcggtggcggagagaacctgtacttc | Used as insert into pACYC-1-GLP-1/2 by In-Fusion |
| 20xGly R | gaagtacaggttctctccgccaccgcctccaccccgccgcctccgcc tccccctccacccccctccaccgcccccaccccccgaagcagc | Used as insert into pACYC-1-GLP-1/2 by In-Fusion |

*Table S 3 - **Primer nucleotide sequence.** All primer sequences used for this study are listed, F, forward; R, reverse.*

| | |
|---|---|
| GLP-1 | 5'-AGGGGGTCAGGTCGAgacgtcAGAGAACCTGTACTTCCAACACTCTGAAGGGACGT TTACTTCCGATGTCAGTTCCTATTTAGAGGGGCAAGCCGCCAAGGAGTTTATTGCTTG GCTGGTAAAAGGGCGTGGTTAGTCGACAAGCTTGCGG-3' |
| GLP-2 | 5'-AGGGGGTCAGGTCGAgacgtcAGAGAATTTATATTTCCAACATTCTGACGGATCATT TAGTGATGAGATGAACACAATCCTTGATAATTTGGCTGCTCGCGATTTCATTAACTGG TTAATACAGACAAAAATTACGGACTAGTCGACAAGCTTGCGG-3' |
| ProcM leader | 5'-TTCGAGCTCGGCGCGaATGTCAGAAGAACAATTGAAGGCTTTCATAGCTAAAGTC CAGGGTGACAGCTCTTTACAAGAGCAGTTAAAGGCGGAAGGAGCTGATGTAGTAGC AATCGCCAAGGCTGCGGGGTTTACGATTAAGCAGCAGGATCTTAACGCGGCCGCCTCAGAGTTATCGG ATGAGGAATTGGAAGCTGCTTCGGGGGGTGAGAACCTGTACTTCCAATAGCGCGCCTGCAGGTCG-3' |
| LctM leader | 5'-TTCGAGCTCGGCGCGaATGAAAGAACAAAATTCTTTCAACCTCCTGCAGGAGGTA ACTGAGTCAGAGCTGGATTTGATTTTAGGGGCTGAGAACCTGTACTTCCAATAGCGCGCCTGCAGGTC G-3' |
| CylM leader | 5'-TTCGAGCTCGGCGCGaATGCTGAATAAGGAAAATCAAGAGAACTATTATTCCAAT AAATTGGAATTAGTAGGGCCGTCTTTTGAGGAACTTAGTTTAGAAGAAATGGAGGCGATCCAGGGCA GCGGTGACGTTCAAGCAGAGGAGAACCTGTACTTCCAATAGCGCGCCTGCAGGTCG-3' |
| HalM1 leader | 5'-TTCGAGCTCGGCGCGaATGACGAATCTTCTGAAGGAATGGAAGATGCCATTGGA GCGCACTCATAACAATTCAAATCCAGCTGGGGACATTTTTCAAGAGCTGGAGGATCAGGACATTTTAGC CGGCGTGAACGGGGCCGAGAACCTGTACTTCCAATAGCGCGCCTGCAGGTCG-3' |
| HalM2 leader | 5'-TTCGAGCTCGGCGCGaATGGTAAATAGTAAGGATCTGCGGAACCCTGAATTCCGT AAGGCACAGGGGCTCCAGTTCGTCGATGAAGTTAATGAGAAGGAACTGTCTAGTTTGGCCGGGAGTG GTGACGTACACGCGCAAGAGAACCTGTACTTCCAATAGCGCGCCTGCAGGTCG-3' |

*Table S4 – Ordered GeneBlocks from IDT. Overhangs in red (for In-Fusion) and TEV site is marked in blue and underlined.*

| | |
|---|---|
| ProcM leader | MSEEQLKAFIAKVQGDSSLQEQLKAEGADVVAIAKAAGFTIKQQDLNAAASELSDEEL EAASGG |
| LctM leader | MKEQNSFNLLQEVTESELDLILGA |
| CylM leader | MLNKENQENYYSNKLELVGPSFEELSLEEMEAIQGSGDVQAE |
| HalM1 leader | MTNLLKEWKMPLERTHNNSNPAGDIFQELEDQDILAGVNGA |
| HalM2 leader | MVNSKDLRNPEFRKAQGLQFVDEVNEKELSSLAGSGDVHAQ |
| NpnM leader | MSEQEQAQTRQDIEARIIAKAWKDEAYKQELVTNPKAVIEREFGVEFPADVNVQVLEENP TSLHFVLPISPVAIAQELSEEQLEAIAGG |

*Table S5 – Amino acid sequence of the leader peptides used in this study.*

| Autoinduction media (Terrific Broth) (g/L) | |
|---|---|
| Tryptone | 12.00 |
| $MgSO_4$ | 0.15 |
| $KH_2PO_4$ | 6.50 |
| Glucose | 0.50 |
| Yeast Extract | 24.00 |
| $(NH_4)_2SO_4$ | 3.30 |
| $Na_2HPO_4$ | 7.10 |
| Alpha Lactose | 2.00 |

*Table S6 – Recipe for autoinduction media.*
